# Volumetric ultrasound localization microscopy of the whole brain microvasculature

**DOI:** 10.1101/2021.09.17.460797

**Authors:** Baptiste Heiles, Arthur Chavignon, Antoine Bergel, Vincent Hingot, Hicham Serroune, David Maresca, Sophie Pezet, Mathieu Pernot, Mickael Tanter, Olivier Couture

## Abstract

Technologies to visualize whole organs across scales *in vivo* are essential for our understanding of biology in health and disease. To date, only *post-mortem* techniques such as perfused computed tomography scanning or optical microscopy of cleared tissues achieve cellular resolution across entire organs and imaging methods with equal performance in living mammalian organs have yet to be developed. Recently, 2D ultrasound localization microscopy has successfully mapped the fine-scale vasculature of various organs down to a 10 *μm* precision. However, reprojection issues and out-of-plane motion prevent complex blood flow quantification and fast volumetric imaging of whole organs. Here, we demonstrate for the first time *in vivo* volumetric ultrasound localization microscopy mapping of the rodent brain vasculature. We developed a complete methodological pipeline that includes specific surgery, a dedicated 3D ultrasound acquisition sequence, localization and tracking algorithms, motion correction and realignment, as well as the post-processing quantification of cerebral blood flow. We illustrate the power of this approach, by mapping the whole rat brain vasculature at a resolution of 12 *μm*, revealing mesoscopic to macroscopic vascular architectures and cerebral blood flows ranging from 1 to 100 *mm/s*. Our results pave the way to the investigation of *in vivo* vascular processes across the mammalian brain in health and disease, in a wide range of contexts and models.

## Main

Blood circulation mediates a fundamental interplay between the extracellular and intracellular space. It carries oxygen, energy, and all fundamental nutrients to cells everywhere in the body. As revealed by William Harvey^1^, blood circulation is a dynamic process that must be studied in the context of a living organism. The vascular system encompasses several scales of vessels, from centimeters wide arteries to capillaries whose diameters are smaller than a single red blood cell. It ensures normal metabolism and function but also drives the progress of the major fatal diseases such as cancer^2^, atherosclerosis^3^, stroke, diabetes^4^, Alzheimers’^5^, or Covid-19^6^.

Comprehensive understanding of normal vascular function, along with disease diagnosis, has led to the creation of imaging modalities capable of mapping blood circulation. In humans, CT, MRI, Ultrasound Doppler^7^, and contrast-enhanced ultrasound^8^ can map large vasculature or evaluate perfusion level in tissue non-invasively. In animal models, finer structures of large portions of the brain can be resolved at the micrometric scale using techniques such as tissue transparization^9–13^, staining ^14^, or micro-CT^15^. They often require equipment that is restricted to pre-clinical use, long acquisition times - from several hours up to several days - but most importantly the micrometric resolution is attainable only *post-mortem*, which precludes investigations of blood flow dynamics. Two-photon imaging in animals through small chronic window implants has imaged the vasculature down to the capillaries at submicrometric scales but are limited to the cortex due to light absorption^16–18^. Photoacoustic microscopy^19,20^ combined with tomography approaches is capable of imaging 3D organs *in vivo*, however, depth of imaging is related to its lateral resolution and so the penetration-resolution trade-off remains.

With the current state of the art, we are faced with the choice between imaging larger vessels over large depths and entire organs or imaging the microvasculature at shallow depth in small regions. Consequently, the microcirculation anatomy and physiology at the scale of an entire living mammalian organ has yet to be observed, a feat vital to measure blood variations induced by neuronal stimulation, follow angiogenesis in tumorous tissue, or reveal the dynamics of the vascular system when submitted to intense stress during strokes or seizures. The need for whole-brain mapping at micrometric scale of animal models *in vivo* is critical to neuroscientists relying on the vascular response to assess neuronal response.

The combined use of clinically approved contrast agents, namely microbubbles, with high frame rate echographs allows us to surpass conventional wavelength-bound resolution. These agents, which remain entirely intravascular, are used as individual ultrasound sources - alike to fluorescent molecules in single-molecule localized optical microscopy^21^-in a technique called Ultrasound Localization Microscopy^22–24^ (ULM). The echo originating from individual microbubbles can not only be localized with a resolution of a few microns ^25^ but can also be tracked over several tens or hundreds of milliseconds, revealing precise perfusion with high temporal sampling. 2D ULM has been applied to various organs such as the brain ^26,27^, the kidney ^28^, and in the context of cancer models to tumors ^29–31^.

However, 2D ULM implementations are inherently limited by the lack of information outside of the imaging plane. Moreover, the projection of ultrasound echoes in one plane makes flow quantification valid only for vessels aligned with the axial direction such as in the neocortex in the brain. Plane limited imaging causes projection artifacts prevents appropriate motion correction and increases user-dependency in the choice of the plane. Finally, while imaging a single plane takes about a minute, a plane by plane approach would involve precise translation and rotation scanning, multiple injections, and prolonged imaging time considerably limiting clinical translation for 3D diagnosis of large and complex structures.

Here, we report a volumetric ULM method relying on a fully populated matrix transducer and custom-built ultrasound scanner that achieves high volume rate 3D imaging at the microscale over an entire living rodent brain. We show that the vascular physiology of the entire organ of the rat can now be characterized with precise super-resolved velocimetry thanks to in-depth optimization of ULM post-processing algorithms to handle the large amount of data produced. We measure this resolution *in vivo* with several metrics taken from the literature and test the robustness of the results with respect to anesthesia. Last, we introduce a clustering method to isolate trajectories into separate entities representing vessels. This adaptive segmentation method allows automated diameter measurement and easier visualization of specific structures directly from millions of microbubble paths. Thanks to these additional post-processing steps, we gain insight into the precise anatomical and hemodynamic characterization of the rodent brain vasculature.

## Results

Several objectives were identified to overcome the challenges presented by 3D ULM such as designing a new experimental pipeline to image the whole brain, extending ULM algorithms in three dimensions and improving them to make computation times realistic while retaining image quality, and exploring the ability of the technique to quantify hemodynamic changes over the whole vasculature in 3D.

### Development of a dedicated animal surgery and a new acquisition and processing pipeline for brain-wide in vivo volumetric ULM

The extension of ULM to 3 dimensions was originally demonstrated in an *in vitro* flow phantom representing a single sub-wavelength bifurcating vessel^32^. The transition to imaging a living organ is complexified by the presence of the skull, increased number of vessels, heterogeneous microbubble speeds, tissue motion, and ultrasound-related problems such as the trade-off between Signal to Noise Ratio (SNR) and volume rate, tissue backscattering, and absorption. We implemented 3D ULM in two phases: in the first animal group, we have imaged the rat brain through a 12×12 *mm* cranial window in a single ULM volume anterior to Bregma^33^. Then, we aimed to image the whole brain of the rat and developed a novel whole skull trepanation surgery, imaged the entire brain with volumes acquired sequentially for different probe positions, and stitched them together with a big-data realignment method. The general framework of ULM is applied to both groups (outlined in black tiles in **figure 1.a**)). Special processing steps are then implemented to either provide fine characterization at vessel scale for the first group or whole-brain hemodynamic imaging (outlined in orange and green tiles on **figure 1.a**)).

**Figure 1.**
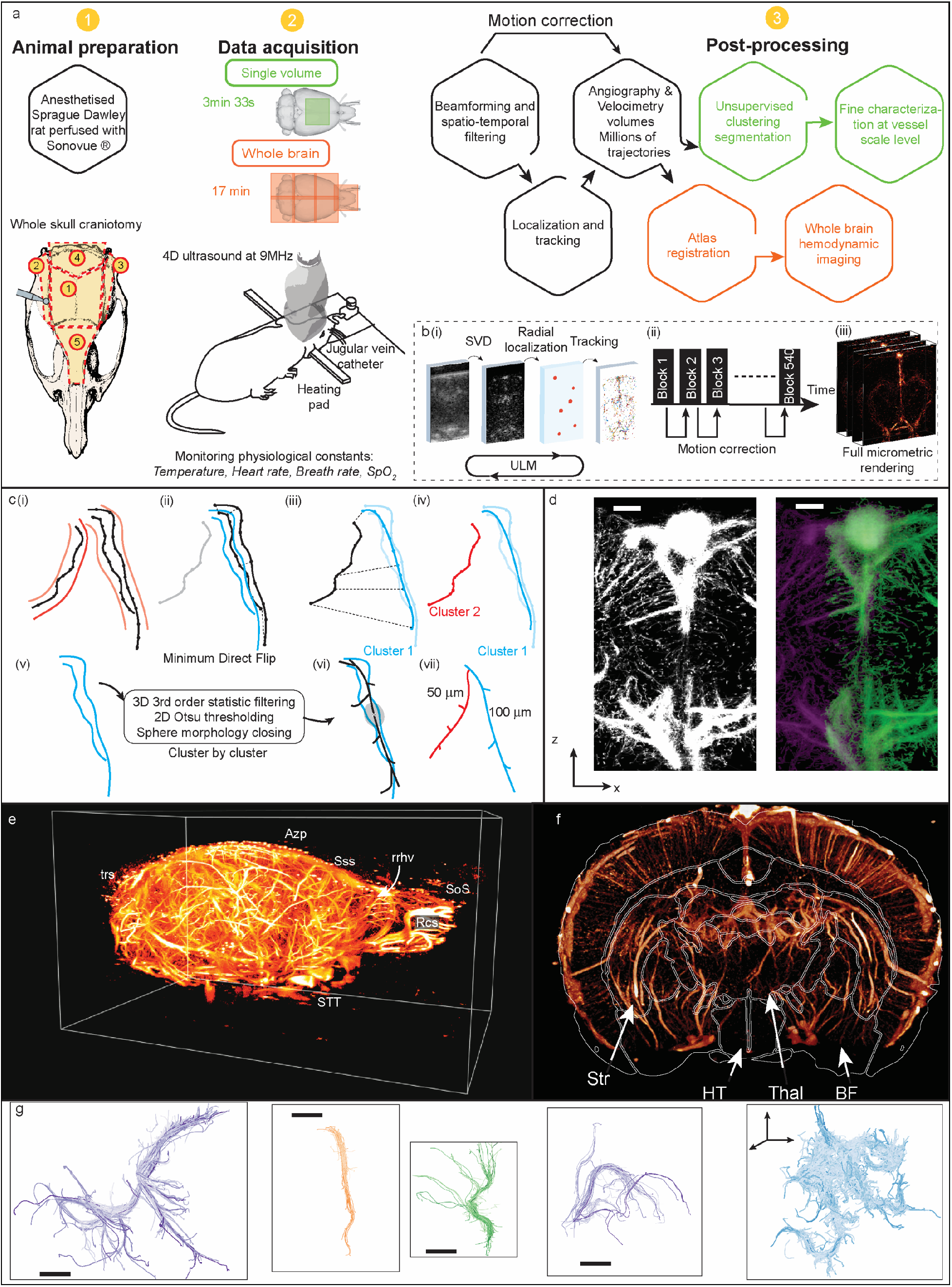
Overview of 3D ULM on the whole brain. a) Presentation of the 3D *in vivo* ULM surgery, acquisition, and post-processing pipeline. The animal is anesthetized and a jugular vein catheter is placed to inject microbubbles. A cranial window is opened: for a single volume (in green in this figure), only area 1 is removed, for whole-brain imaging, the 5 windows are removed sequentially indicated by circle numbers. Data acquisition is done with 4D ultrasound at 9MHz central frequency and the animal is under constant physiological monitoring. In post-processing, the data is then beamformed, filtered, localized and trajectories are reconstructed. Post-processing steps are added for single volume (in green) to perform fine characterization at vessel level and for whole-brain imaging (in orange) to stitch volumes and register with a Waxhom space-based atlas b) 3D ULM specific processing: i) show filtering, localization and tracking applied to all volumes acquired. ii) presents the motion correction algorithm in between noncontiguous volumes. All the volumes are acquired in a block like fashion, with a pause in between each block. A rigid transformation is calculated on tissue signal from the first volume of each block (Block 1, Block 2,…). Then this transformation is applied directly to the trajectory points. iii) the motion corrected trajectories are used to produce density or velocity maps. c) Segmentation processing illustrated on two neighboring vessels. i) The supposed borders of the vessels are drawn in red. Assuming we want to analyze 3 trajectories. ii) First, the Minimum Direct Flip distance is calculated between the two blue trajectories. As their MDF value is below threshold, they are paired and make up a new cluster defined by its centroid streamline. iii) Second, the MDF from the other trajectory to the centroid streamline is calculated. iv) Because it is above threshold, it is used to create a new cluster. v) For each of the clusters, the velocity volume is rendered and pre-processed with 3D 3^rd^ order statistic filtering, Otsu threshold to yield a binary image which is then morphologically closed. vi)This volume is then used for medial axis thinning skeletonization, pruned and an ellipsoid fit is used to calculated diameters vii). d) Stitching of two volumes along the superior sagittal sinus: Maximum Intensity Projection on the left and Green Cyan combination on the right. (scale bar 200 *μm*)). e) Whole organ imaging of a Sprague-Dawley rat brain: 3D rendering showing the right part of the hemisphere. Azp: azygos pericallosal artery, ios: inferior olfactory sinus, Rcs, Rostral confluence of olfactory sinuses, rrhv: rostral rhinal vein, Sos: Superior olfactory sinus, STT: spinal trigeminal tract, trs: transverse sinus (retrogloid vein) f) Coronal view (Antero-posterior axis Bregma −0.8 mm) and sagittal view (Medio-lateral Bregma +0 mm). Str: striatum, HT: hypothalamus region, Thal: thalamus, BF: Basal Forebrain region. Overlaid with Waxholm space atlas ^36^ g) Clusters from different regions rendered from the microbubble trajectories: From left to right, 4 clusters make up an artery in the striatum (light purple), one cluster representing a cortical vessel (orange), one cluster representing the thalamoperforating artery (green) and one cluster sufficient for the rendering of the transverse hippocampal vein (purple) (scale bar 500 *μm*). Finally, 27 clusters render the complex vascular structure descending from the longitudinal hippocampal vein down towards the septal region in 3D. The color intensity encodes the length of the trajectories.

The sequence was optimized to reach a higher Signal-to-Noise Ratio, essential *in vivo*. Due to higher speckle and tissue noise than in the *in vitro* phantom, we had to increase the number of compounded angles to 12^34^, also increasing the data size. The general ULM processing is illustrated in **figure 1.b) (i)**: after delay and sum beamforming, the volumes were filtered with Singular Value Decomposition to remove tissue signal in ultrasound images and improve the detection of microbubbles in 3D. To cope with lower SNR and with increased data size, we designed a new 3D localization algorithm based on the radial symmetry of the Point Spread Function (PSF). This new localization approach has been shown to retain high imaging quality while cutting processing costs by a factor as much as 50 when translated in 2D^35^. In parallel, the beamformed data were filtered to retain only tissue signal and used for motion correction in between the different blocks (**figure 1.b) (ii)**). The rigid transformation calculated in that step was then applied to the corresponding trajectories. Finally, density and velocity rendering are obtained by compounding all the trajectories (**figure 1.b) (iii)**). This whole process was parallelized to keep calculation costs to four hours.

To perform whole-brain imaging, we imaged 7 ULM volumes at different positions mapping the whole brain (**figure 1.a**)) with an overlap of at least 200 *μm*. This was performed under anesthesia after careful removal of the skull bone while leaving the dura mater intact. The total imaging time is 17 minutes. The total size of the dataset – half a million independent frames, and the 7 reconstructed ULM volumes-is several terabytes big. The realignment of the 7 different volumes was streamlined to the CPU processor using a rigid transformation. This operation was applied using the C *elastix* toolbox and relied on pyramidal multi-resolution down-sampling of the volumes to keep computational costs to a minimum. The volumes were then registered with the Waxholm Space atlas for Sprague Dawley rats^36^ and the whole brain was rendered with density and velocity compositions. A cross-section of one millimeter is shown in **figure 1.d**). On the left, a Maximum Intensity Projection centered on the superior sagittal sinus is rendered. The stitching is not visible. For comparison, a composed image is shown on the right with cyan and green reflecting the colors of the two volumes. The sinus is well stitched and so are the smaller cortex vessels plunging downwards, regardless of their orientation.

For hemodynamics analysis, we devised a segmentation and clustering pipeline based on trajectories rather than individual voxels. We first devised a pre-processing step to avoid the heavy computational cost of processing the whole volume at once and proceed vessel by vessel. We regroup all *N* trajectories in *M* clusters before implementing segmentation and skeletonization. This approach was inspired by a similar one used in Diffusion Tensor Imaging (DTI) and fiber tracking^37^. Further detail is given in the methods section, but the complexity of the clustering was reduced from *O*(2*N*)*^N^* to *O*(*MN*), *M* « *N*. As illustrated in (i), this technique allows to combine trajectories that have the same shape and are closely located together in structures. At initialization, the first cluster is taken as the longest trajectory available. To create the other clusters, the Minimum Direct Flip distance from each trajectory to all of the existing cluster centerlines is calculated (**figure 1.c**) (ii)). Based on a threshold, this trajectory is either added to the cluster (iii)) or used to create another cluster (iv). Because the number of trajectories for each ULM volume is around a million, the reduction of complexity is crucial. Then, each cluster is reconstructed into a volume that is further filtered to yield a binary image. Skeletonization with pruning (v), and diameter measurement via ellipse fitting (vi) is then carried out. This post-processing step added after ULM enables us to perform organ-wide hemodynamic analysis in an unsupervised fashion (**figure 1.a**) green tiles).

### *3D imaging of the whole rat brain* in vivo *below the diffraction limit*

As seen in **Figure 1e**), the implementation of 3D ULM results in maps of the vascular network of the rat brain. The volumetric image presented here has its voxel encoded with an intensity relating to the number of microbubbles tracks passing through that pixel. The microbubbles being intravascular contrast agents, this intensity can be seen as an indirect measure of local blood flow. The whole-brain ULM displays the olfactory bulb with its preeminent inferior and superior olfactory sinus down to the cerebellum with its median medullary arteries^38^. The top of the cortex is split by the superior sagittal sinus into which several veins drain. On the pial surface of the brain, we can observe the different branches from the anterior, middle, and posterior cerebral arteries. Deeper structures are visible like the azygos pericallosal artery, and below the hemispheres, the spinal trigeminal tract as well as the optic nerves.

In **Figure 1 f**), we rendered a coronal slice of 400 *μm* of the whole brain. The vasculature is clearly visible, especially in the cortex where hundreds of penetrating arteries are observable. The complex vascular tree in the striatum and fibria of the hippocampus is also visible. Deeper, within the midbrain, arteries ascending from the Circle of Willis can be seen although the density of the vasculature renders precise identification delicate. Hundreds of penetrating arteries are observable within the cerebral cortex. We have registered the whole brain with the Waxholm space Atlas^36,39,40^ by using regional landmarks common to the vascular and tissue structures. Additional coronal, transverse and sagittal views are available in **Supplementary Figure 3**. This enables us to segment anatomical zones and their vascular content but also to compare several organs together. These structures form the basis of a 3D vascular atlas that can be exploited to identify each part of the brain.

Finally, thanks to the unsupervised clustering method developed here, it is possible to visualize vessels independently by rendering their trajectories. This allows easier navigation along the millions of trajectories and easier classification of the different vessels in the functional zones defined in the atlas. In **Figure 1**Figure *1* **g**), we have represented a few interesting structures from different regions in the brain. Depending on the size of the vessel, one cluster can represent all of it – in light purple, an artery in the striatum, orange, a single cortical vessel, in green the thalamoperforating artery – but aggregating more clusters together allows us to look at whole areas, for example, 27 clusters in blue render the vascular structure descending from the longitudinal hippocampal vein.

### Resolution measurements

The resolution of the technique was measured using three indices from the state of the art in the single volume experimental paradigm. The anesthesia was also changed to investigate the robustness of the resolution measurements in between volatile Isoflurane (Iso) and injected Ketamin-Medetomidine (KM). The approach we followed was to separate one ULM volume into 8 isotropic sub-volumes of size 5.5 × 5.5 × 6 *mm* (**Figure 2a**)) by halving each direction once, providing statistical measurements of resolution and allowing to reconstruct volumes on finer grids without computational constraints.

**Figure 2.**
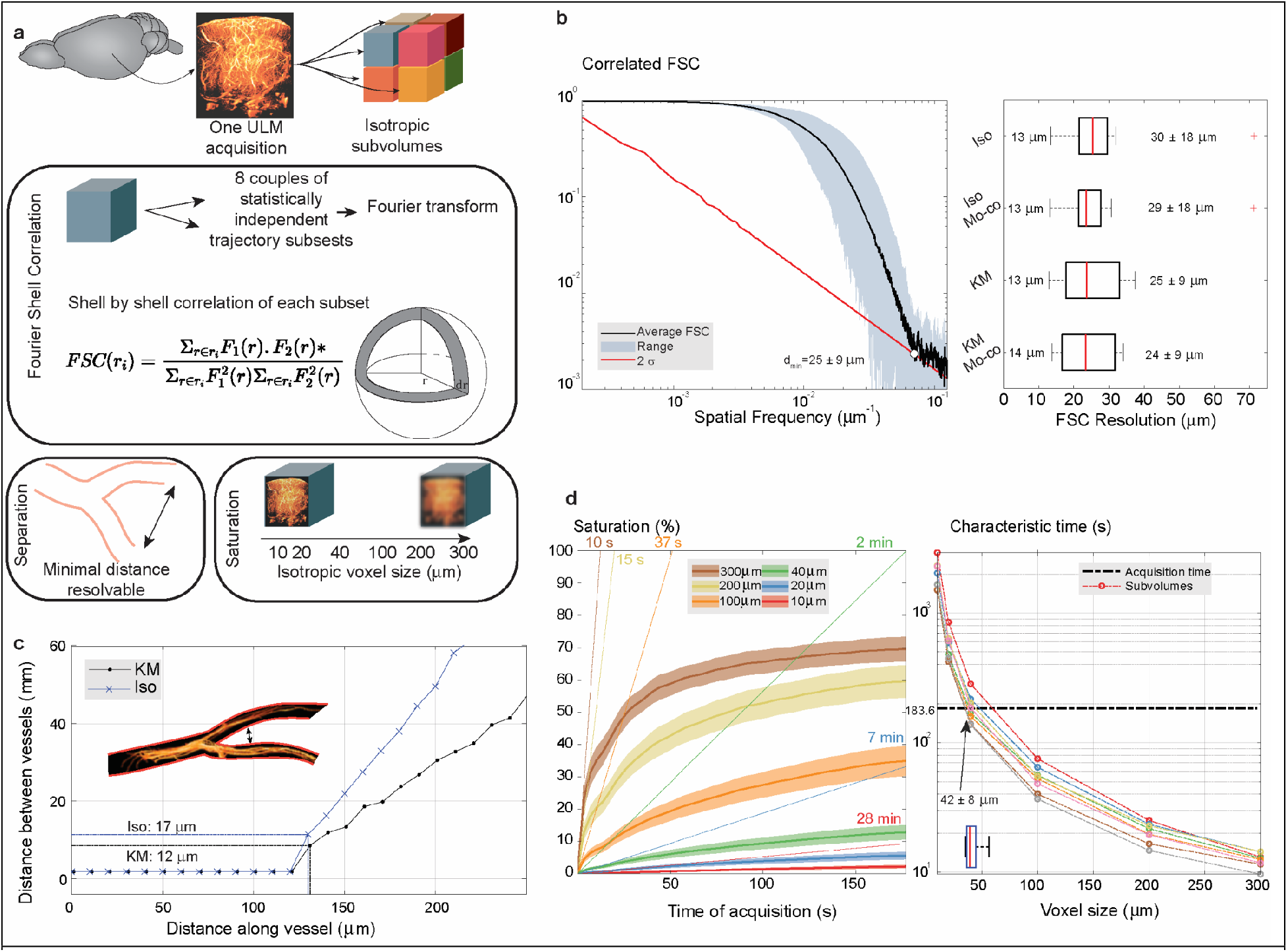
Quantifying resolution on a single ULM volume with 3 state of the art metrics. a) Illustration of the three different approaches used to quantify resolution: First, a single volume ULM density rendering is divided into 8 isotropic sub-volumes. These sub-volumes will be used to calculate the resolutions b) The range of the FSC is presented in blue, overlaid with its average in solid black and the 2*σ* curve in red. This resolution is calculated for one ULM volume acquired under both anesthetics (Iso/KM) and with/without motion correction. Values are represented in a boxplot with whiskers indicate upper and lower quartiles. The minimum value is on the left of the boxplot while the average and standard deviation are written on the right c) Separation index: the distance between the two main branches of the striate artery clustered below is calculated, the minimum non-zero value indicates the minimum resolution attainable. This index is calculated for both anesthetics Iso and KM. d) Resolution measured with saturation: in color, saturation for different voxel sizes with respect to time is measured (average is overlaid in solid line over standard deviation in transparent curves). Next, the characteristic time calculated from saturation is plotted in function of voxel size for each of the 8 sub-volumes. The acquisition time is indicated in a dotted black line.

The first index explored was the Fourier Shell Correlation^41,42^ a measure that computes resolution from a threshold of correlation in between independent subsets of the volume. For each of the 8 sub-volumes, the trajectories were separated in two statistically independent groups and a volume was reconstructed with a 4 *μm* isotropic voxel. This volume was then transformed in the Fourier domain and, the logarithmic cross-correlation coefficient over corresponding shells in Fourier space is represented for a ULM volume. The mean curve was overlaid with the standard deviation for each spatial frequency value in gray shade (**figure 2.b**)). The 2*σ* curve is plotted and the resolution is found as the first intersection with the cross correlation^43^. On average, the resolution was determined to be 29 ± 17 *μm* with a minimum value of 13 *μm* reached for one of the sub-volumes. The same was done for the same rat with a different anesthetic, the mean resolution was found to be 24 ± 9 *μm* and a minimum of 12 *μm* was reached, showing no statistical difference (paired t-test, *p* = 0.49). The resolution measurements were calculated for motion corrected datasets (Iso Mo-co/ KM Mo-co).

A second way to measure resolution is to come back to its literal definition and consider the ability of the system to separate neighboring structures^32^. The trajectories in these arteries were reconstructed on an isotropic grid with voxel size 10 *μm* and the vessels were aligned with their centerline’s principal axis (see Methods). The distance between the two branching canals was measured between the two closest trajectory points from the outside borders of the canals. The minimal distance measurable in the ISO case was 17 *μm* 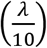 (**figure 2.c**)). The same process was repeated for the KM case and the distance was found to be 12 *μm* (λ/14).

The 8 sub-volumes were also analyzed for their resolution with respect to saturation^33^. After being reconstructed with isotropic voxel sizes of [300,200,100,40,20,10] *μm*, the saturation time was calculated and plotted as a function of pixel size. The total time of acquisition used to reconstruct this ULM volume is 3 min and 33 s and is plotted as a red dashed line (**figure 2.d**)). On average the theoretical resolution reachable for this time of acquisition is found to be 4 times better than the wavelength at 9 *MHz*, at 42 *μm*, with a minimum of 27 *μm* (*λ*/6). The worst-case scenario was also sub-wavelength at 75 *μm*. The results for the same rat with a different anesthesia are also presented and show similar results (*mean* = 65 *μm*, *min* = 32 *μm*, *max* = 80 *μm*).

### Case study: the striate arteries

To showcase our ability to image deep into the brain we have chosen to illustrate a number of concepts and possibilities of ULM in a specific case. An interesting functional region is the striatum. Prone to stroke but also involved in reaching mechanisms, Parkinson’s disease, and addiction^44–46^, we will focus on the irrigating arteries of this region to expose hemodynamic quantification at the micrometric scale. They branch out from the middle cerebral artery (**Figure 3.a**)) in successive structures with decreasing diameters arranged in a lattice-like fashion^47^.

**Figure 3.**
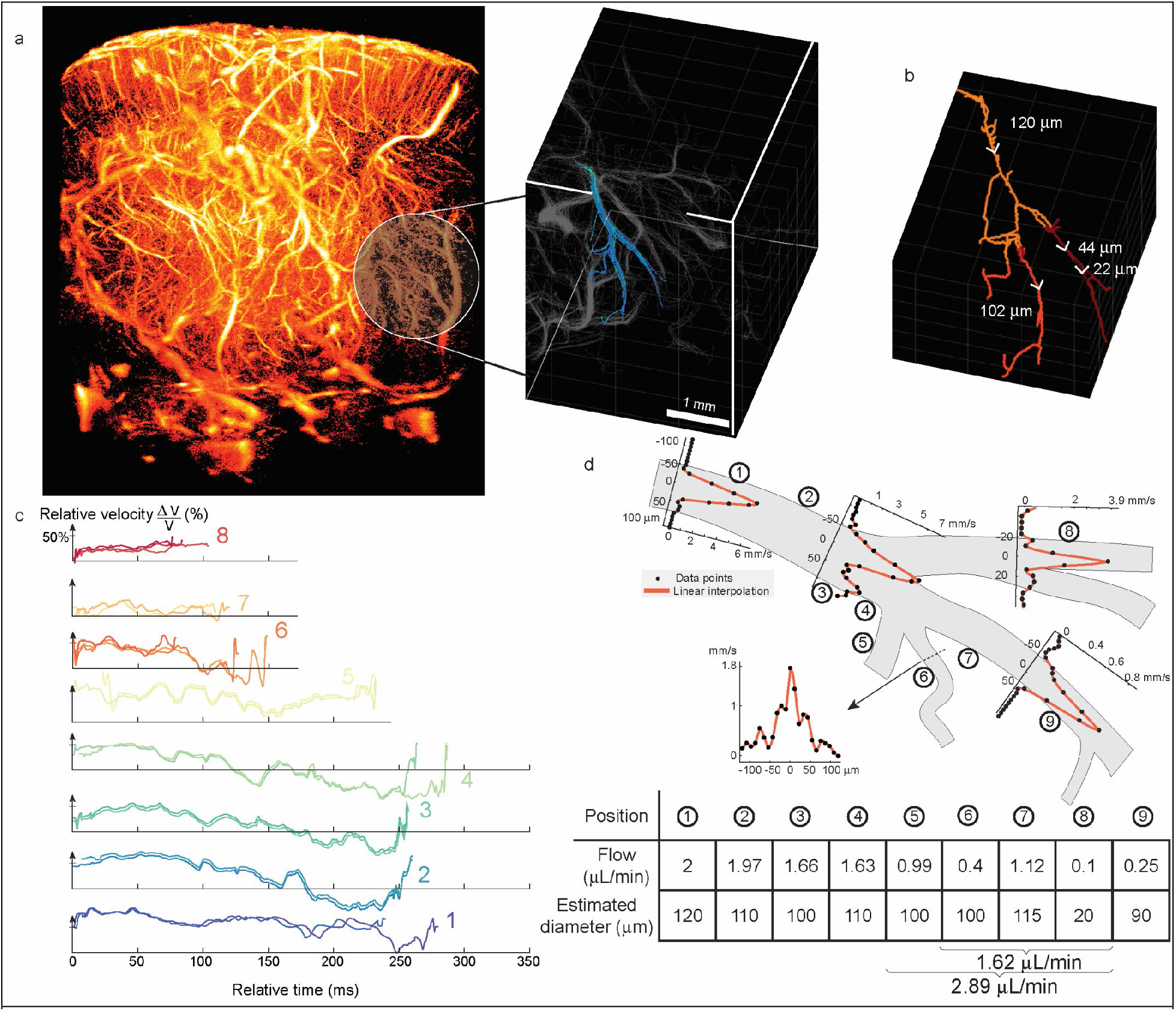
Case study: Striate arteries. a) ULM volume of the rat brain from an offside coronal view: Intensity corresponds to the density of microbubbles passing through each voxel. The zoomed-in portion on the right shows trajectory-based representation. The trajectories not belonging to the cluster including the striate arteries are in gray, while the ones of interest are color-coded according to the length of the trajectories. The motion was corrected in between each block with rigid transformation. b) Measurement of diameters inside the striate arteries: the diameters were calculated as the short axes’ dimension of an ellipsoid fitted along the centerline containing all trajectories. The average diameter between two nodes after segmentation is indicated here with a color code. c) Velocity dynamics of microbubbles inside 17 trajectories: a few trajectories inside the striate artery have been selected and their velocity variation 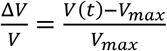 are represented in function of time. These trajectories were grouped together based on their velocity variation over time. The velocity variation inside trajectories are representative of the pulsatile flow present in the arteries. As such, microbubbles in the same vessels and acquired during the same heartbeat exhibit the same patterns. Because of discontinuous block-to-block acquisition, some trajectories are interrupted mid-acquisition, which explains why they don’t span the same time frame. d) Wall to wall flow profiles in the striate artery and flow analysis: The outer border of the realigned projected striate artery is represented in black. For 5 different positions, the flow profiles are taken as the average velocities over a 840 *μm* ~5*λ* slab colinear to the maximum flow streamline. The velocity points are represented in solid black dots and a spline interpolation model in solid orange curve. The axes are indicated in micrometers and mm/s. Circled numbers indicate where the flows and diameters are calculated in the table.

The whole volume was clustered and we identified the anterior striate artery as a single cluster (represented with length of trajectory color in **Figure 3a**), overlaid on the other clusters represented in gray). The volume presented here measures 3 × 5.5 × 7mm^3^ and is located 4 *mm* deep below the cortex surface and is reconstructed with a voxel size of 10 × 10 × 10 *μm^3^*.

The cluster was skeletonized and the diameter of each vessel was measured at every point of the skeleton. The average diameter per segment of the skeleton is represented in **Figure 3 b**). The decreasing size of the upper branch of the artery is apparent in both our measurement and the trajectory-based image. The smallest diameter measured was 22 *μm* while the main branch of the vessel measured 120 *μm* on average.

Thanks to the high temporal sampling of ultrafast ultrasound, we can track the velocity dynamics of each microbubble trajectory. **Figure 3 c**) presents plots of trajectories from individual microbubbles in function of time and color coded according to their length. The relative variation of speed was found to be as much as 94 % in the longest trajectory. Pulsatile flow is observable in most trajectories where the speed is modulated by heartbeat and packets of trajectories exhibit similar peak-to-baseline variation. The shorter the trajectories are, for example 6; 7; 8, the less they exhibit variation of velocity but their dynamics remain similar. The heartbeat of the rat during this acquisition was measured to be around 300 bpm on average, yielding a heart cycle of 0.2 *s*.The longest trajectory we observe here lasts for 0.28 *s*, which is consistent with the observation of a single pulse in the velocity dynamics.

In **Figure 3d**), we have outlined the vessel walls and superimposed the velocity profiles calculated at several points. These profiles are derived from a spatially averaged velocity calculated over 840 *μm* (5*λ*) in the direction of the flow and aligned with the main axis of the vessel. The microbubble velocity in this vessel reached a maximum of 7.2 *mm/s* and could be as low as 1.4 *mm/s*. At the branching sites (3 and 4), one can see the velocity profiles getting separated upstream and exhibiting two distinct maxima in their distribution. Profiles taken a few hundreds of micrometers downstream of the bifurcation return to fully developed profiles. At each bifurcation, the profiles show the same behavior, separating in two, with higher velocity towards the main branch. We have calculated the flow assuming a Poiseuille profile and the diameter based on the profiles (see Methods). The total flow upstream from the branch at points 6, 7, and 8 gives us a value of 1.62 *μL/min*, while the original flow closest to the beginning of the branch is evaluated at 1.66 *μL/min*, in good agreement with conservation of flow. The values calculated with the flow are on par with the values calculated in the skeletonization.

## Discussion

We have demonstrated 3D *in vivo* imaging of the vasculature below the diffraction limit. Based on ULM which was originally demonstrated in 2D the physical principle stays the same but the extension to 3D has required extensive rework of the animal model as well as a redesign of the post-processing techniques. To perform 3D ULM in an acceptable time, the acquisition method, as well as the image formation algorithms, were changed to adapt localization and integrate motion correction schemes. By adding the third dimension, ULM is now able to accurately depict the vasculature’s orientation and tortuosity, as well as measure velocities without suffering from re-projection. We were capable of mapping structures as close as 12 *μm* apart, render the cerebral vascular system with a minimum FSC based resolution a tenth of the wavelength at 17 *μm*, and capturing hemodynamic changes deep within the living brain. Finally, by applying this imaging technique sequentially after complete skull removal, the whole brain was rendered with a 12 *μm* voxel grid along with blood velocities and registered with a Waxholm based atlas. Complemented by trajectory-based clustering and flow analysis algorithms, this enables ULM to leap forward as a quantitative imaging tool for vascular applications.

Brain-wide imaging with 3D ULM, which can be applied to all organs, has the tremendous advantage of visualizing the vascular dynamics at play over a wide range of spatial scales. Sampling all microbubbles at the same time over large areas allowed to depict complex structures and to follow their trajectories regardless of the vessels’ orientation, which is out of reach of current techniques. The advantage of such an accurate representation of the vessel lies in the ability to calculate diameters precisely which could then be used to propagate flow models inside each vessel. Ideally, imaging microbubbles in 3D rather than in 2D also allows reducing acquisition time per injection as all microbubbles contribute to image construction rather than just the ones in the field of view. For the same volume of imaging, 3D ULM is thus much faster than plane by plane ULM, and only requires one bolus. As shown in **Figure 2 c**), if we were to base the measurement of resolution on the saturation, the resolution we found by calculating the characteristic time is on average 42 *μm*, one order of magnitude larger than what is found in 2D at an ultrasound frequency of 15 *MHz* (Hingot et al ^33^ calculated around 5 *μm* for a 4 minute acquisition). However, this approach postulates that all pixels should be saturated, which is not always the case regardless of the dimension. The lower resolution we obtain compared to the 2D case is to be put in perspective as the frequency of the probe is different and the 4D ultrasound scanner is an innovative design and not a commercial one. Higher frequency fully populated^48^ or sparse array probes are beginning to emerge as valid alternatives to fully addressed probes and we expect our technique to be translatable to any probe design^49,50^.

The comparison of the different resolution criteria and their results when calculated on the same animal undergoing different types of anesthesia, vouches for the robustness of 3D ULM. The non-significance of the statistical test for the Fourier Shell Correlation based resolutions presented in **Figure 2b**), shows that regardless of the physiological condition during the recording, the final resolution of the image stayed the same. Because the final images are composed of millions of trajectories taken at different times of the cardiac cycle, they tend to integrate the variation in frequency content that is brought by hemodynamic changes. However, as highlighted in Figure 3 Case study: Striate arteries, such changes can still be picked up in individual microbubble trajectories. This demonstrates that while 3D ultrasound localization microscopy in its final super-resolved volume of the whole organ is a minute-long process, it is composed of individual events sampled at very high temporal resolution, giving access (though partially) to physiologically induced variations locally. The technique is thus a combination between a global approach where the organ’s vascular anatomy can be represented with micrometric precision but every few minutes and a local approach where the organ’s hemodynamic activity is described on the millisecond timescale at sparse spatial coordinates. If one is interested in a specific region in the brain, multiple trajectories can be extracted and dynamic information about cerebral blood flow can be recovered. The characterization of flow outlined in **Figure 3 d**) is made possible thanks to the information of velocity and an accurate representation of the vessel in the three dimensions. However, we see that conservation of flow is not respected when we take into account vessel 5 into our calculations (the flow is higher by 45% at 2.89 *μL/min* instead of 2*μL/min*). We hypothesize that vessel 5 is irrigated by another artery that we have not considered into the clustering method. Because the clustering algorithm is sensitive to the order the trajectories are presented in^37^, here it may be failing because the longest trajectories are used first to build clusters and then smaller trajectories are paired with the clusters. As such, the algorithm does not pair large vessels together but pairs smaller vessels to the larger ones. The speed of this algorithm outweighs greatly its ability to pair clusters together.

Our study remains limited in several aspects. The technique presented here was applied to a trepanned rat brain. The animal model was adapted to maximize the SNR of the images and to be able to deliver the highest resolution possible. Transcranial ultrasound localization microscopy is possible ^27,52,53^, but the aberration of the ultrasound waves, when it goes through the skull, is likely to result in a decreased imaging quality. Interesting algorithms to correct bone aberration have been put forward. The brain was chosen to demonstrate the possibility of the technique because its motion can be limited through the use of a stereotactic frame. Our method can be applied to any organ but would require partially adequate motion compensation schemes such as the ones presented here. For larger motions, imaging quality can benefit from applying volume to volume correction rather than block by block.

Due to technological limitations such as the size of the acquisition data, we accumulated a number of microbubbles insufficient to characterize the capillary bed^33^. Methods to increase the number of detectable microbubbles^51^ could be exploited along with compressed-sensing approaches, to drive the cost of acquisition to a minimum. Several approaches have been developed in 2D that might be of interest to 3D should they be extended. Deep learning methods to find microbubbles buried deep down in the non-filtered data should also be of interest but the cost in memory of developing these to 3D remains quite high. The time it takes to acquire the 3D ULM representation of the brain also constitutes a limitation to our technique as the physiological variations of the whole organ will not be measured at the same moment. We only possess information where microbubbles are located at a precise point in time and so can represent hemodynamic changes over limited scales. In our implementation, the heart rate, breathing rate, temperature were controlled throughout the whole experiment, and anesthesia, heating pad temperature, oxygen inflow were adapted to limit the alteration of the physiological constants.

Our probe was limited in size/number of elements and, in the case of rats, the whole-brain imaging needed realignment of several acquisitions. While the anatomical description of the organ is scarcely affected by this thanks to sub-voxel precision of the realignment, the hemodynamics are not as microbubble trajectories are cut-off in between volumes. However, due to the semi-continuous imaging of the block-to-block approach, microbubble trajectories are already usually cut off. The emergence of continuous imaging in 3D with large probes for long acquisition times by using sparsely addressed probes looks promising to be able to fully sample organs through space and time continuously. Finally, this stitching approach has to be put in perspective to what is done in 2D optical scanning imaging techniques, where every plane needs to be registered.

The results presented here have required extensive post-processing. The steps related to localization, tracking, and filtering are relatively easy to get accustomed to, on the other hand, the clustering, segmentation, and hemodynamics processes are much more complicated and finely tuned for our application. The unsupervised fashion in which the clustering method operates already simplifies the workflow by just needing to adjust the Minimal Direct Flip value and two or three thresholds. To apply our technique to other organs, adaptation of these algorithms and methods, changes might be needed. We foresee that other approaches for example based on machine learning-based will emerge to have a more generalized method of hemodynamic characterization, similarly to what has recently been published in the field of tissue clearing^12,13^.

Finally, the results shown here depict the anatomical and hemodynamic features of the brain but do not have access to biological or chemical markers. The use of tagged microbubbles with proteins to monitor biomarkers has been brought forward in the field of biomolecular ultrasound^57^. The combined use of such contrast agents will give access to biomarker mediated ultrasound super-resolution similarly to what has been shown in clearing techniques^10^ and in optical superresolution^58^. The application of brain-wide ultrasound localization microscopy as a pre-imaging step for other applications such as functional ultrasound is also of great interest. Other organs could also benefit from *in vivo* micrometric imaging such as the heart and the small vasculature spurring from the coronaries, the intestines, where the vasculature decreases to micron sizes and is responsible for key nutrient transport, or the renal cortex where the interglobular arteries supply the glomeruli, the beginning point of the kidney’s filtration system. While all these applications are of tremendous interest, 3D ULM comes short of increasing both spatial and temporal sampling at the same time, which confines it to the post-imaging analysis of the hemodynamics. It would be beneficial to restore the higher temporal sampling brought by the use of high frame rate ultrasound and breach the gap between functional imaging, which necessitates a temporal resolution smaller than a second, and super-resolution. With further improvement in matrix transducer and portable scanners, volumetric ULM has the potential to be a tool for the longitudinal study of the microvasculature of entire organs, along with its variations for physiological or functional studies. Development in smaller and more targeted contrast agents^59–62^ could extend its applications to the extravascular space.

## Conclusion

We report volumetric ULM, a vascular super-resolution ultrasound imaging technique surpassing the classical resolution limit by a factor larger than 10. We used 3D ULM to fully image a trepanned rat brain in vivo, successfully captured the entire cerebral vasculature, and co-registered it with a Waxholm space-based atlas. In each artery and vein depicted, we could derive blood velocity by calculating the velocity of ultrasound contrast agents circulating in each voxel. This enabled us to map hemodynamic vessel parameters in 3D. We foresee many applications for this technique in the future ranging from anatomical description of the vascular system to hemodynamics-based diagnosis of stroke, tumor microenvironments, or any deep investigation of microvascular function in opaque tissues.

## Material and Methods

### Animals

All animals received humane care in compliance with the European Union Directive of 2010 (2010/63/EU), and the study was approved by the institutional and regional committees for animal care (Comité d’Ethique pour l’Expérimentation Animale no. 59—‘Paris Centre et Sud’ Protocole no. 2017-23).

8-10 weeks Sprague-Dawley rats were obtained from Janvier Labs (Le Genest-Saint-Isle, France). Animals were free from pathogens and were placed in our in-house facilities. Housed in Techniplast ^®^ cages by groups of 2 minimum and 3 when the total animal per volume ratio did not exceed values fixed by regulations, the animals were kept for at least a week before surgery. The temperature in the cages was controlled to be between 21-23°C, the humidity rate in between 40-60%. The day-night cycle was divided as follows: 7h-19h/19h-7h. Water and a commercial pelleted diet SAFE A04-10 were available ad libitum. SAFE Flake sawdust was used as bedding, enrichment such as pieces of cardboard, paper tunnels were provided and the facility vet went to visit them daily. This allowed to reduce stress during manipulation before anesthesia induction and increase the anesthesia’s performance in terms of speed and robustness in time.

After 5 minutes in the induction cage filled with a mix of air and 5% of Isoflurane, the animal was placed on the back on a heated plate with a respiratory mask. The mix was replaced with O2 mixed with 4% of Isoflurane. The depth of the anesthesia was tested by the absence of withdrawal reflex when pinching the toes of the hind limb. An incision was made just above the first rib in the thoracic area. The absence of movement from the animal during that incision was considered as a confirmation of the depth of the anesthesia. After the tissue was dilacerated to provide easy access to the jugular vein, two suturing strings were threaded under it. The string closer to the brain was knotted to close blood flow returning from the brain. An incision perpendicular to the direction of the vein was made and a 180 *μm* inner diameter polyethylene tubing (Instech/A-M Systems ^®^) was placed inside the vein to serve as a catheter. The surgery was deemed successful when blood reflow was observed. The skin was sutured back with veterinary glue (Vetbond ^®^ 3M Animal Care Products, St-Paul, MN, USA).

The animal was then placed in a stereotactic frame equipped with a respiratory mask. The percentage of Isoflurane was decreased to 3% while maintaining the absence of reflexes. 5mL of saline at 37°C was injected subcutaneously in two different doses on the left and right sides of the back of the rat to prevent dehydration. Such a procedure was repeated every 2 hours. The fur was cut off the scalp using an animal hair trimmer to provide a clear view of the skin and a clean surgery site. The skin was cleaned first with a 4% povidone-iodine scrub solution, and second with a 10% povidone-iodine solution for additional antiseptic action (both from (Betadine^®^, Purdue Products, LP, Stamford, CT)). An incision was made in the scalp following a line between Bregma and Lambda points but extending from over the olfactory bulb to just behind the cerebellum (see **Supplementary** Error! Reference source not found.**1 step 1, 2, 3**). The scalp was retracted and the side and the cerebellum muscles were disjoined from the skull to allow easy access to the skull. The muscles were kept away from the skull by using suturing thread. After this, the percentage of Isoflurane was decreased as low as possible but above 1% to prevent lethal side-effects such as hyperventilation, cardiac deficiencies while maintaining the absence of reflexes.

A drill was used to remove a large portion of the skull to provide an ultrasound transparent window. This window was drilled using 5 different zones (see the layout on **Supplementary** Error! Reference source not found.**1 step 4**). The first zone is the biggest one starting over the lambda sutures, went along the sides of the skull just below the ridges, and then joined 3 mm after the Bregma point just before the beginning of the olfactory bulb. Such a window measured approximately 12 mm in width and 15 mm in height. This window was extended to the sides and then two additional zones were drilled to provide access to the cerebellum and the olfactory bulb. The surgery took between 2 and 3 hours. A saline solution was used to keep the dura mater hydrated at all times.

### Physiological constants

The heartbeat, breath rate, breath distension, temperature, and SpO_2_ were all monitored using MouseOx Plus pulse oximeter (Starr ^®^ Life Sciences Corporation). A decrease in SpO_2_ was remedied by an increase in the O_2_ fraction of the gas mix breathed by the animal. A decrease in temperature implied covering the back of the animal with a heated glove until the appropriate temperature was recovered. An increase in temperature implied lowering the temperature of the heating plate. A sudden decrease in breath rate was usually solved by lowering the percentage of Isoflurane in the mix. Breath distension measurements were discarded because of a large standard deviation due to inaccuracies in the measurement device.

### Isoflurane and Ketamin-Medetomidin acquisitions

For the anesthesia paradigm experiments, only zone 1 was removed. The surgery took in between 30 and 45 minutes. The imaging process began 30 minutes after the skull was removed. In the meantime, the brain was immersed with saline which will also serve as an ultrasound coupling medium. This allowed the physiological constants to stabilize and be less affected by surgery. We recorded these and considered them as a baseline for the rest of the experiment. Just before the injection of Ketamine-Medetomidin, Isoflurane was decreased to around 0.8% and then decreased by 0.2% steps in less than one minute after the injection of Ketamine-Medetomidin (KM). This allowed preventing overdosing the animal. After Isoflurane was stopped completely, we waited for 60 minutes to make sure that the Isoflurane completely disappeared in the blood and that its effects were null after that time. Additional time was sometimes required to make sure the physiological constants returned to the baseline. KM was injected every hour and a half after the first injection or if reflexes were observed. A half dosage was injected to prevent overdosing.

### Ultrasound sequence

We designed our ultrasound sequence around the computational capabilities of our system. In our case, working with RF data is mandatory for the 3D approach as the 1024 channel ultrasound scanner is unable to beamform as it is acquiring data.

A customized, programmable, 1024-channel ultrasound system ^63^ was used to drive a 32-by-35 matrix-array probe centered at 8 MHz with a 90% bandwidth at −6 dB, a 0.3-mm pitch, and a 0.3-mm element size (Vermon, Tours, France). The 9th, 17th, and 25th lines of that probe are not connected resulting in a 32×32 matrix array. We used that probe to transmit 2-D tilted plane waves at 9MHz frequency at 12V. The probe was coupled to the brain using either water, ultrasound gel, or low concentration agar (less than 2% in water). It was placed directly over the brain with a spacing higher than 3 mm to avoid the near field. The ultrasound system consists of four different Aixplorer synchronized systems (Supersonic Imagine, Aix-en-Provence, France). A combination of 12 tilted plane waves was transmitted with this order: *xz*(°) = [−3, −2, −1,1,2,3,0,0,0,0,0,0]; *yz*(°) = [0, 0,0, 0, 0, 0, −3, −2, −1,1,2,3], with *x* and *y*-directions parallel to the matrix-array probe. We increase the angles to match the maximum number of transmits allowed by the systems transfer rate without dropping the compounded volume rate below 500 *Hz*. The voltage was increased as much as possible to allow higher signal to noise ration while preventing microbubble destruction. The peak-negative pressure (PNP) is −294 *kPa*, and the ISPTA is 385 *mW/cm^2^*. To prevent data overload and to succeed in prolonged high frame rate imaging, a specific sequence was developed with only a few milliseconds of imaging and a long pausing time: during 0.370 s, 185 compounded volumes were acquired with parameters above. The data was then transferred to Solid State Drive memory on each of the four ultrasound scanners’ computers. In total, 540 blocks of 185 volumes were acquired during 18 minutes (see **Supplementary** Error! Reference source not found.). To provide continuous perfusion with microbubbles, 0.1 mL boli were renewed every 90 seconds, yielding a total volume injected of 1.2 mL in 18 minutes which is well below the recommended value of 5mL/kg for a single bolus recommended by the Institutional Animal Care and Use Committee (IACUC).

After acquiring all of the data, it was entirely transferred to another computer where storage has a larger capacity. This data was then processed according to the processing described below.

### ULM processing

The ULM processing is based on the radial symmetry algorithm. The radiofrequency data is beamformed in all directions on a 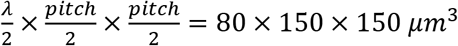 grid. The singular value decomposition is done on each of the 185 volumes ^64,65^. Of the 185 singular vectors, the first 4 are discarded and the volume is reconstructed using 181 of the lowest energy vectors. The radial symmetry-based localization algorithm^66^ is used for localization with an assumed full width at half maximum of 5 pixels in the three directions. The positions of the microbubbles are then converted from pixels to micrometers. The tracking algorithm is implemented on each block of acquisition separately. Tracks are interpolated by resampling the streamline on an oversampled time vector, this way saving the velocity information. Velocities are calculated using differentiation on a custom-built time vector corresponding to the interpolation of the trajectories. The results of this process are the list of non-interpolated microbubble centers, the list of trajectories with the velocities measured at each point of the trajectories. The beamformed data is not saved in the computer’s hard drives. The parallel implementation of this process takes one and a half minutes for one block if velocimetry is performed, only one minute if no velocimetry is performed, making the whole implementation for 540 blocks of 1TB run in 4 hours on a high-performance desktop computer (Intel Core i9 @ 2.9 GHz 12 cores, NVidia RTX 2080Ti, 128GB RAM @ 2133 MHz).

After processing, all trajectories were binned on voxels of size 10 *μm* except when resolution assessment was done. The intensity of each voxel reflects the number of trajectories crossing this voxel. For velocity renderings, the velocity in each voxel is the time average value of the velocities of each microbubble passing through that voxel. For trajectory-based representations, the velocity at each temporal and spatial point is taken.

### Segmentation

To characterize each vessel imaged with 3D ULM, a segmentation algorithm was used. The issue we face was that the conventional segmentation algorithm used in vascular imaging rely on continuous representation of vessels. In our case, vessels are not represented continuously but by the trajectories of microbubbles passing through them. That hinders the robustness and success of the majority of pre-filters used in intensity-based skeletonization algorithms relying on 2^nd^ order derivatives of image intensity, which in our case is not a continuous function inside vessels. Furthermore, each volume generates around a million trajectories, a number so high that conventional algorithm based on independent streamlines (such as the ones used in Diffusion Tensor Imaging (DTI)^67^ or in trajectory analysis for road control^68^) will fail to produce a result in acceptable calculation time.

We decided to rely on a pre-processing step on trajectories where we regroup all trajectories in what we call clusters before implementing segmentation and skeletonization. This approach was inspired by a similar one used in DTI and fiber tracking^37^. All trajectories are first sorted by descending length. For each trajectory, the trajectory is discretized on a cubic spline curve and resampled to yield a reduced number of voxel points *N_reduced_* = 40. Each track thus will have the same number of coordinates and the minimum direct flip (MDF) distance can be calculated. This distance is defined as the minimum of the distance between the corresponding points of two trajectories from start to end, and the distance between the corresponding points of one trajectory to the flipped version of the other. If that distance is smaller than a fixed threshold *MDF_thresh_* = 80, then the two streamlines are paired together to form a new cluster. Should the algorithm compare all tracks sequentially, the number of operations would be (2*N*)*^N^*. In our implementation, a trajectory is compared to existing clusters, and if the minimum MDF value from the trajectory to all clusters is below a fixed threshold, then the trajectory is added to the concerned cluster. Otherwise, a new cluster is created containing that trajectory. When a trajectory is added to a cluster, we calculate a cloud of points belonging to all trajectories in the cluster by concatenating all trajectories in a single one and discretizing it in *N_reduced_* coordinates. These cloud coordinates will be used to calculate the MDF value from the cluster to the trajectories, dictating whether the cluster should be updated with the trajectory or should a new cluster be created. This differs from the centroid approach used in fiber tracking where the centroid streamline is calculated for all trajectories in a cluster. The problem with the centroid method is that it is highly affected by inhomogeneities in track lengths as shown in^37^. In our case, all trajectories belonging to a vessel have different lengths, depending on where they are in the vessel: longer ones are in the middle of the vessel’s lumen because they belong to fast-moving microbubbles and smaller ones are close to the vessel’s boundary as the microbubbles are slowed down by friction from the walls.

After clustering, for each cluster, a volume is built from the velocity measurement. Its squared intensity undergoes 3^rd^ order statistic filtering in 3D on 26 neighbors, followed by binarization at 30% level of the Otsu threshold in 2D and morphological sphere closing of radius 3 voxels. Then, a skeleton is built via a medial axis thinning algorithm followed by branch pruning with a minimum length of 50 *μm*. For each point in the resulting skeleton, the diameter is calculated as the small axes dimension of an ellipsoid centered at this point containing all trajectories inside its volume.

### Measuring velocity profiles from trajectories

The trajectories in a cluster or a volume can be reconstructed on an isotropic grid with voxel size 10 *μm*. On this volume, 3 successive Radon transforms were applied to calculate the principal axis for rotation. Each rotation was applied to the trajectory coordinates to align them on a plane containing most of the trajectories. This allowed us to consider the main direction of flow to be straightened out in the direction 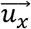 We could then measure diameter, flow, and speed by simply summing along specific dimensions. To do this, used flow profiles and took the first points from the center of the profile (maximum value of velocity) that were less than 20%. Diameters were calculated along the two main dimensions, and the final diameter was taken as the average of the two, 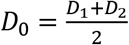. For the flow, we hypothesized a fully laminar profile and took the average velocity to be 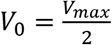. Then the flow was simply taken as *Q*_0_ = *V*_0_π*R*_1_*R*_2_.

### Fourier Shell Correlation

The trajectories belonging to each of the 8 sub-volumes were separated into two subsets randomly. For each subset, the sub-volumes were reconstructed with each voxel having a value representing the number of interpolated trajectories passing there. The size of each voxel was 3 *μm* in each direction. Such a low value was chosen to be able to find the intersection between 2*σ* curves and the Fourier Shell Correlation (FSC) values. The Fourier transform was calculated using the `fftn` function in Matlab. Then, for each Fourier shell, the FSC was calculated as the correlation between the two random subsets. Unfortunately, because of RAM constraints, lower voxel values were not possible. Finally, the FSC was calculated for each spatial frequency along with the *σ*, 2*σ*, 3*σ* curves and plotted.

### Saturation

Saturation^33^ can be calculated for each subset of trajectories in 3 dimensions by reconstructing volumes with different voxel sizes. For each subset, we discretized the trajectories on a pre-defined voxel size matching the resolution to investigate We chose to investigate 7 different resolutions ranging from the pitch value of the probe (300 *μm*) to the chosen voxel size for rendering (10 *μm*): [10; 20; 40; 100; 200; 300] *μm*. A voxel is considered to be complete as soon as a trajectory has passed through it. The total number of voxel was calculated and divided by the subset volume size to yield the saturation value. The time to saturation was measured as the inverse of the tangent at the origin of the saturation curve.

### Separation measurement

To measure the separation in the striatum, we concentrate on the branching present in that vessel. The striatum artery was segmented thanks to the clustering method. The trajectories in the cluster were reconstructed on an isotropic grid with voxel size 10 *μm* and 3 successive Radon transforms were applied to calculate the principal axis for rotation. Each rotation was applied to the trajectory coordinates to align them on a plane containing most of the trajectories on the x-axis. This allowed us to consider the main direction of flow to be straightened out in the direction 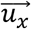. We then measured the distance between the two branching canals by slicing through the vessel orthogonally to the direction 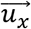 using the L2-norm between the two closest trajectory points.

