## Supplemental figure1 for "Volumetric ultrasound localization microscopy of the whole brain microvasculature"

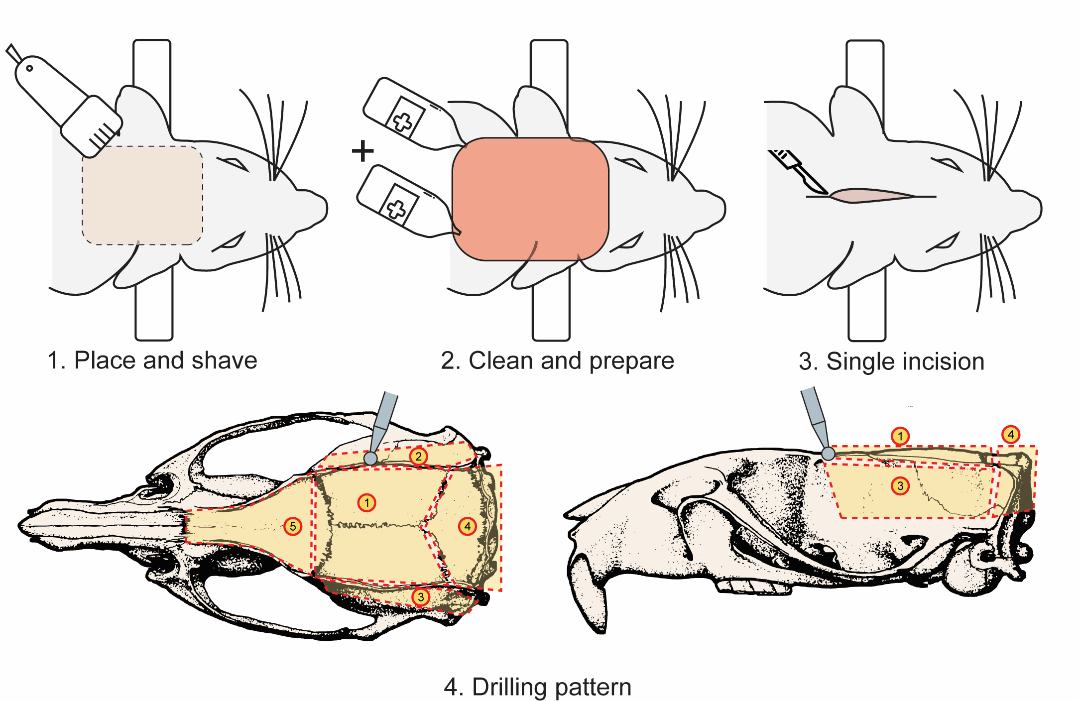

Supplementary figure 1. Steps to remove the whole skull from the brain

1. After the jugular vein catheter procedure, the rat is placed in the stereotactic frame using ear bars. A large zone of fur is trimmed (outlined with a dashed line).
2. The skin on top of the skull is cleaned twice with 4% and 10% povidone-iodine solution
3. An incision is made in the skin and the muscles are retracted from the skull. The surgery site is maintained open with suturing threads.
4. Schematic of the drilling patterns (outlined with dashed line). Zone 1 is the biggest but the easiest to remove as the skull sutures are followed and drilling is made just below the side ridges where the skull is thinnest. Zone 4 just above the cerebellum is prone to bleeding as such it is done directly after zone 1. Then it is left to recover while drilling zone 5 over the olfactory bulb.

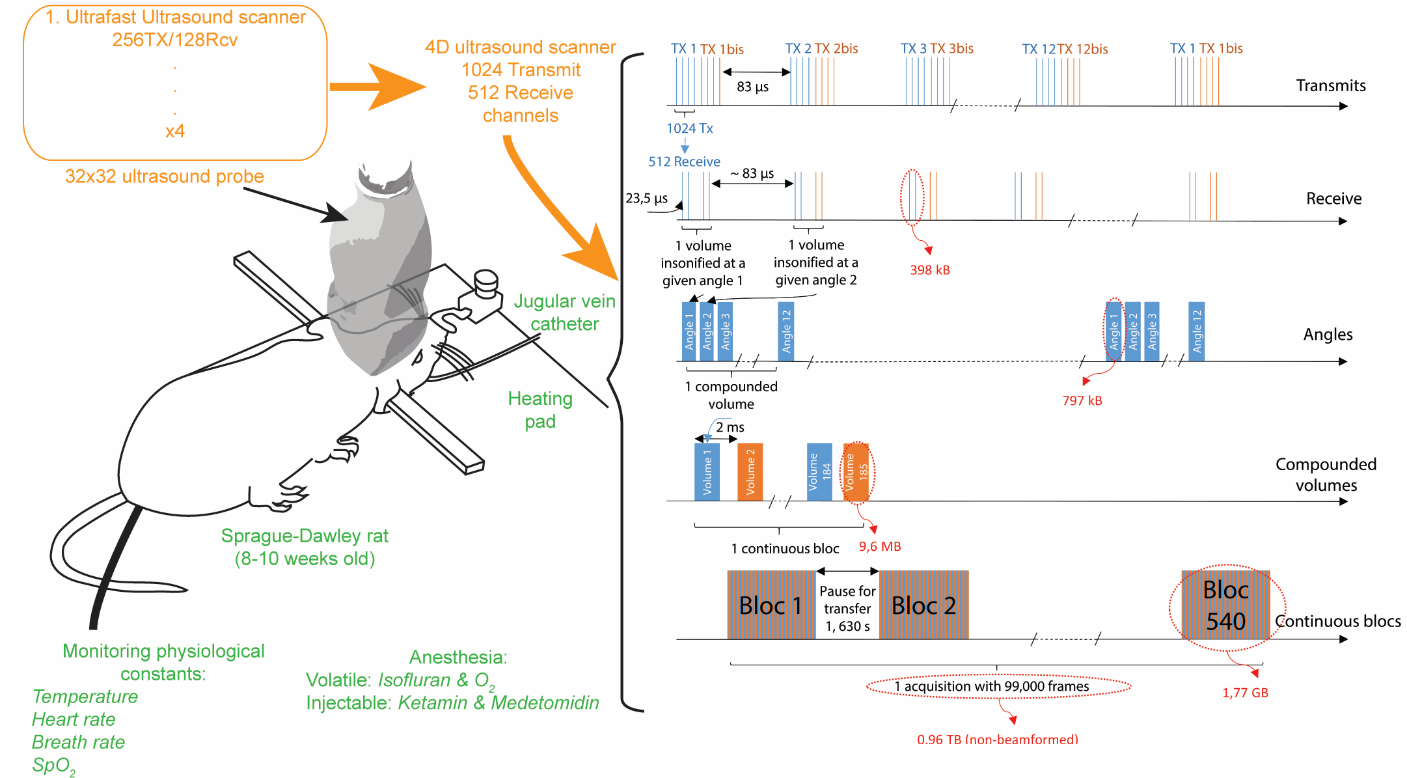

Supplementary figure 2. Setup and ultrasound sequence

The 32x32 fully populated matrix probe is connected through 4 connectors to a custom-built 4D ultrasound machine. This machine is capable of driving 1024 elements in transmit and has 512 receiving channels, meaning each transmit has to be doubled per receive. On the right of the graph, the transmit and receive pattern are drawn. The blue and orange pattern denote first and second set of transmit. We use 24 transmits to make up 12 angles for each volume. A compounded volume is repeated every 2 ms to reach 500Hz volume rate. To allow the data to be transferred to the hard drives, a block-to-block pattern is implemented with a pause in between each transfer of 1.630 s. The whole data for one single ULM volume weighs 0.96TB.

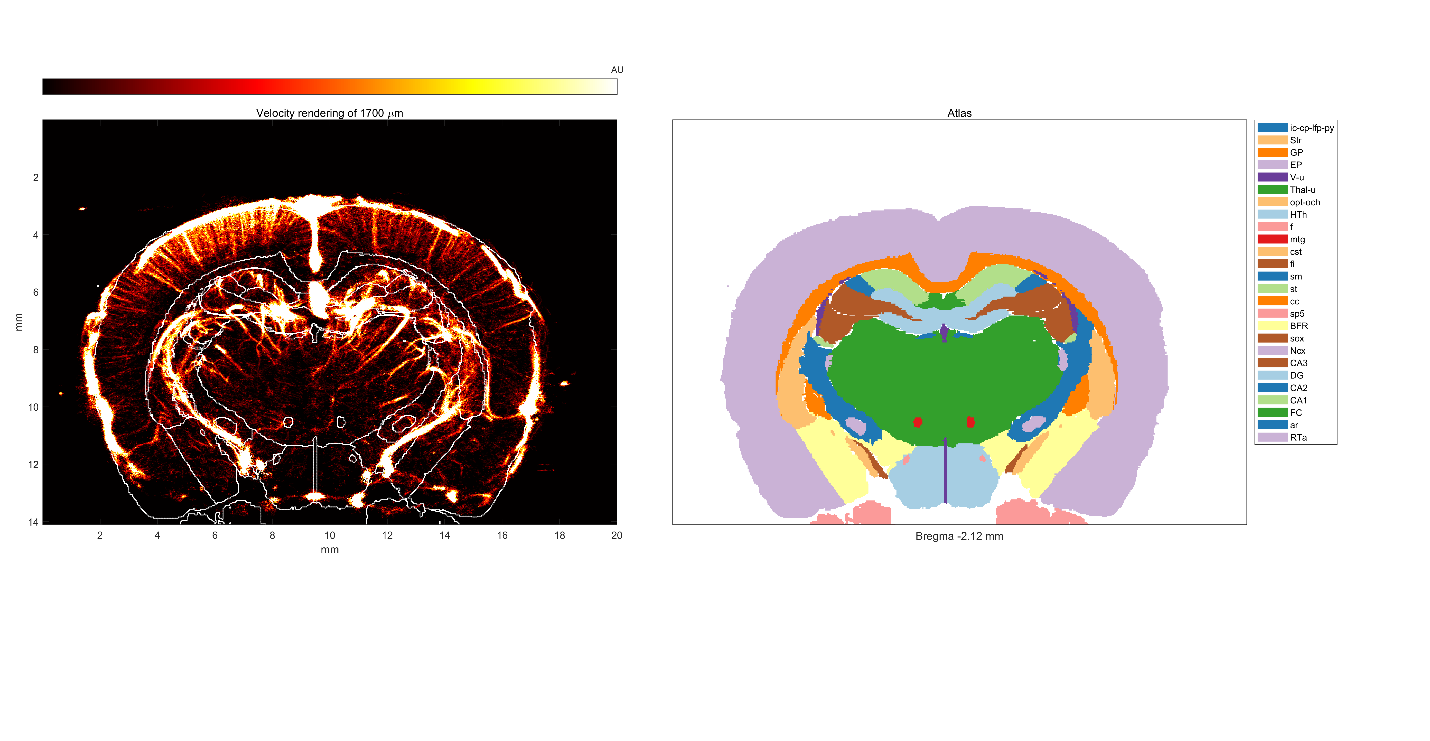

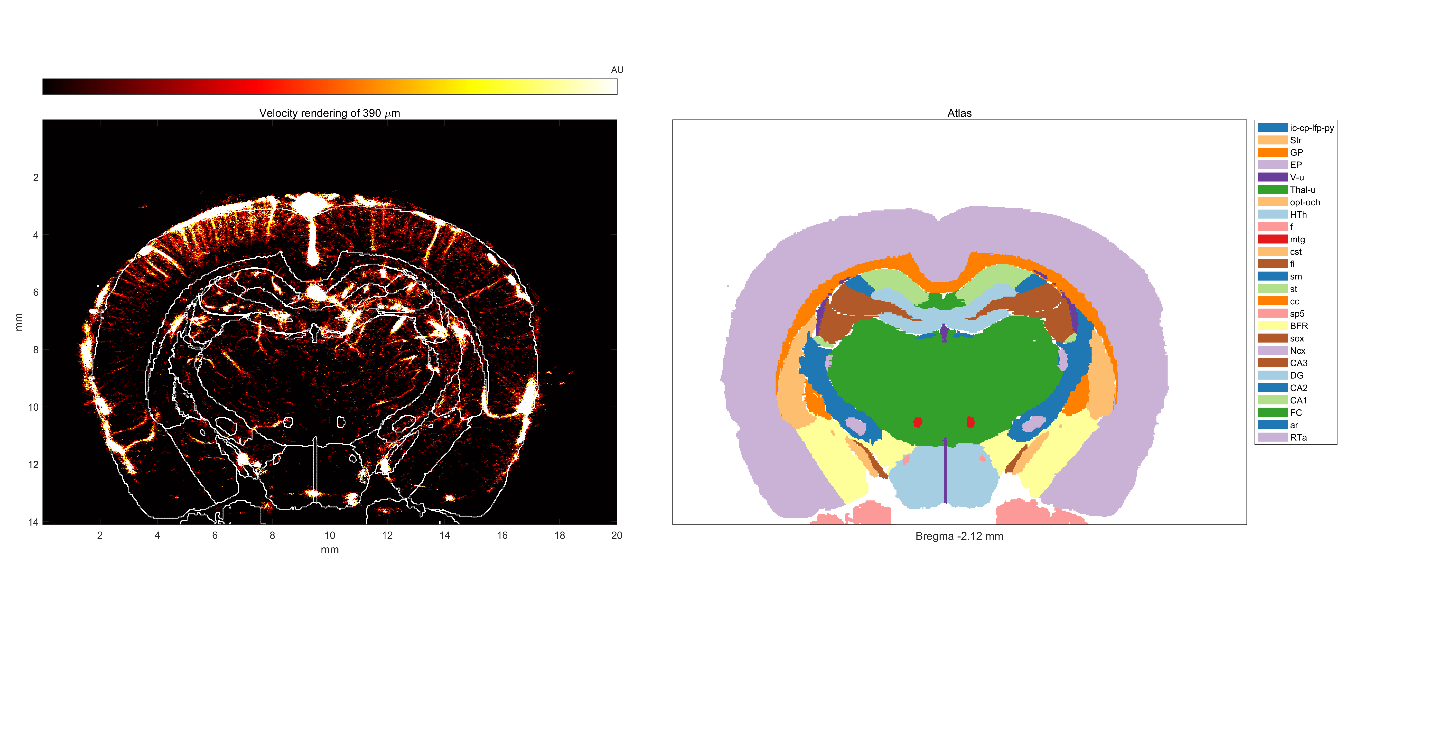

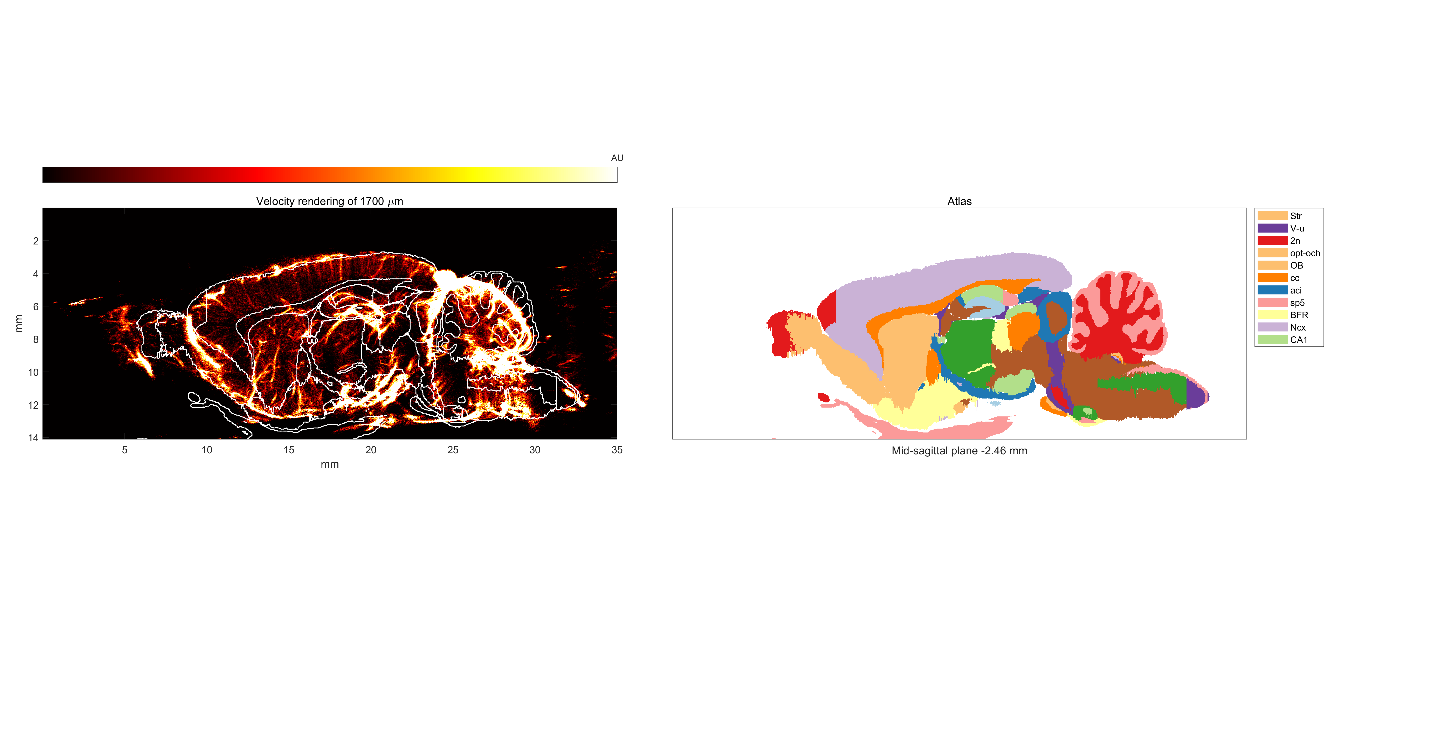

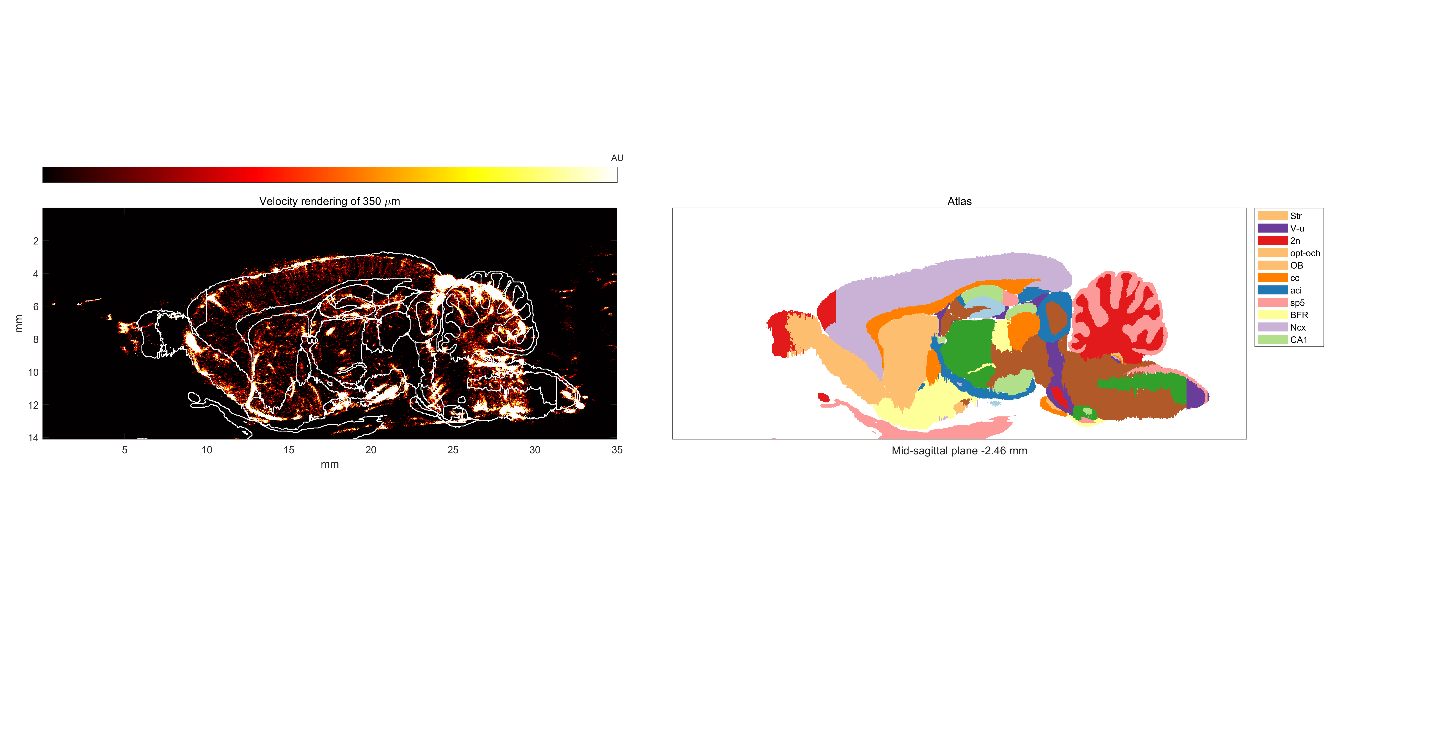

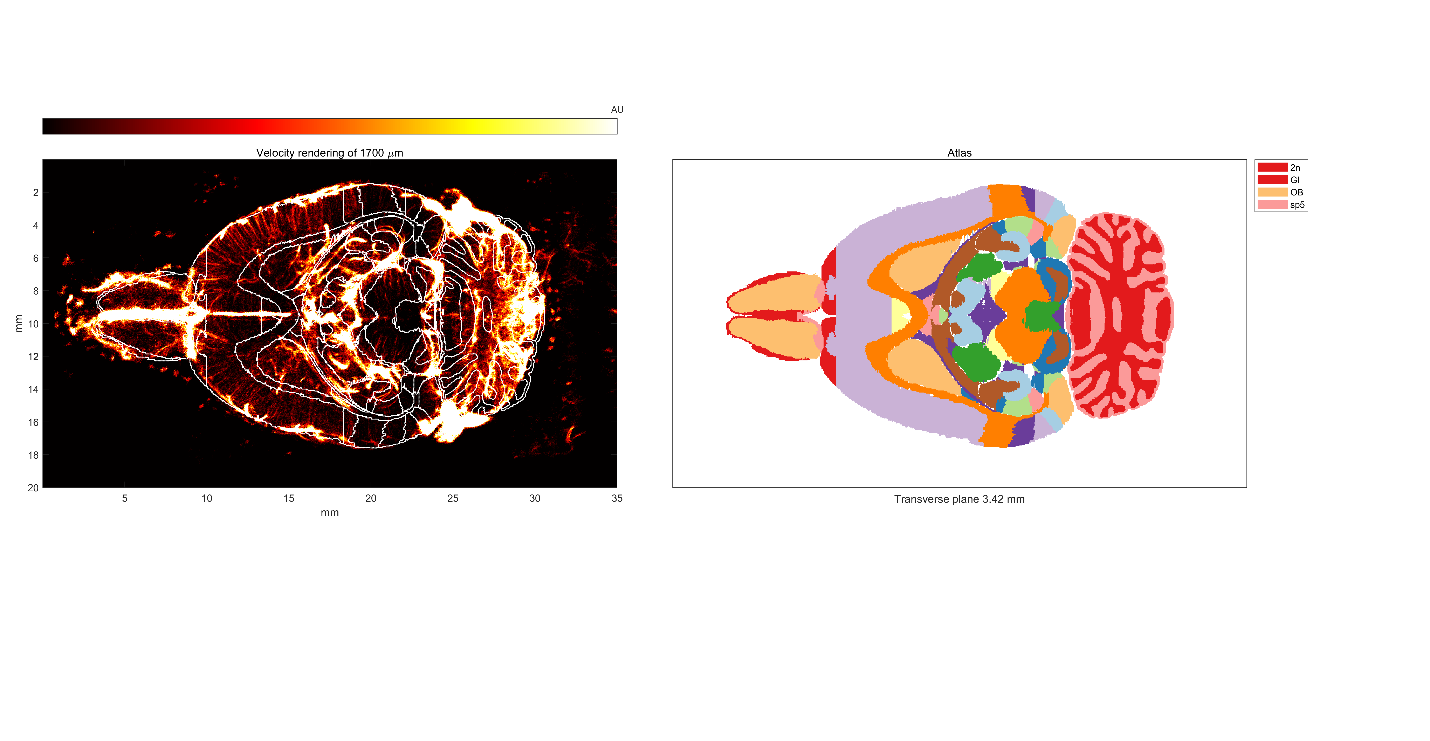

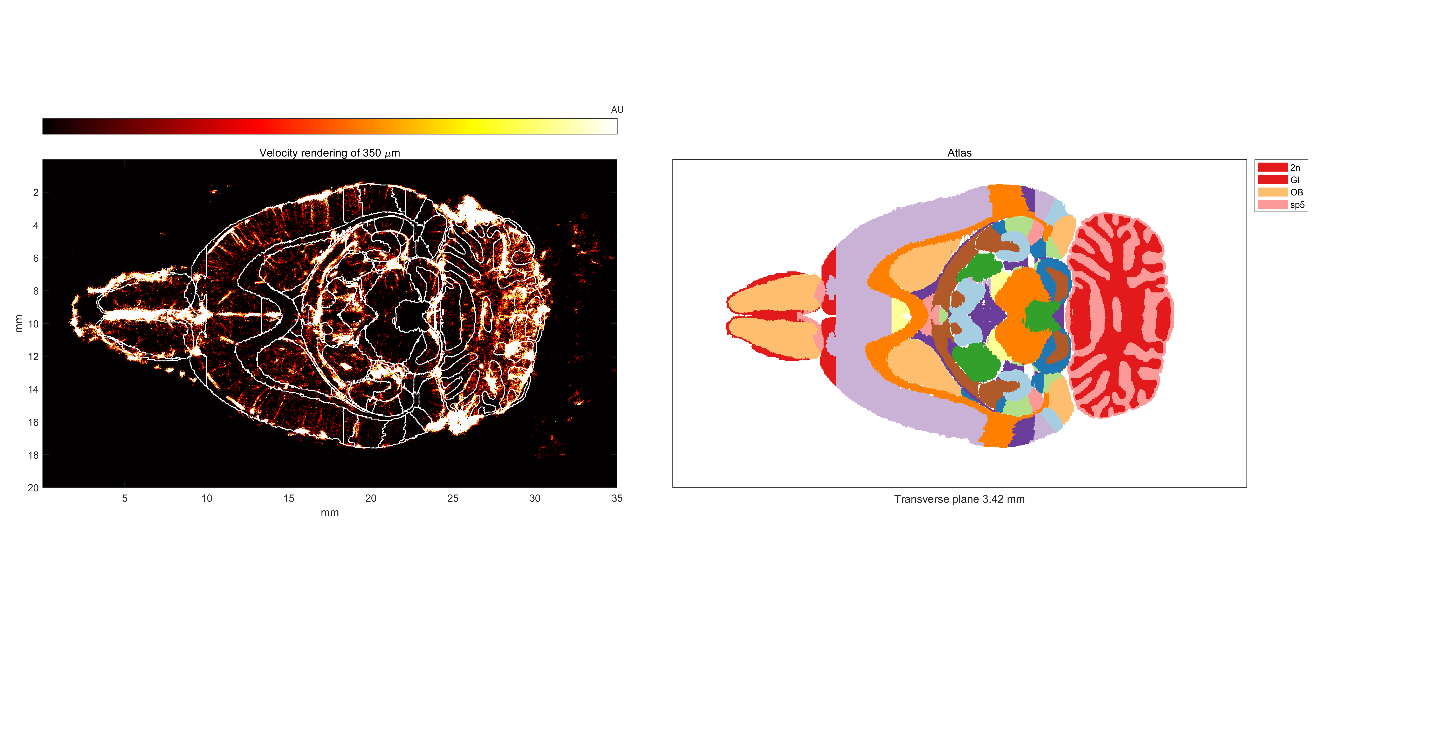

Supplementary figure 3 Volumetric ULM renderings registered with Waxholm-space atlas

Coronal, sagittal and transverse views of 2 slices with different thicknesses corresponding to $10\lambda=1.7 mm$ and $390 \mu m$ which matches the size of 10 slices of the atlas used. ULM images are reconstructed from 7 ULM volumes stitched together and manually registered on Waxholm-space atlas. Abbreviations indicate functional areas as delimited in (Sergejeva et al., 2015)
